# Polarization gratings aided common-path Hilbert holotomography for high-throughput lipid droplets content assay

**DOI:** 10.1101/2024.06.14.598984

**Authors:** Piotr Zdańkowski, Julianna Winnik, Mikołaj Rogalski, Marcin J. Marzejon, Emilia Wdowiak, Wioleta Dudka, Michał Józwik, Maciej Trusiak

**Affiliations:** Warsaw University of Technology, Institute of Micromechanics and Photonics, Boboli 8, 02-525 Warsaw, Poland; Structural and Computational Biology Unit, European Molecular Biology Laboratory (EMBL), Heidelberg, Germany

**Author notes:** These authors contributed equally: Piotr Zdańkowski, Julianna Winnik.

## Abstract

In this contribution we present a novel polarization gratings aided common-path Hilbert holotomography (CP-HHT) for high-throughput 3D refractive index imaging. Addressing limitations in current holotomography methods, we leverage the extended space-bandwidth product (SBP) through robust phase demodulation using Hilbert spiral transform. Thanks to the application of polarization diffraction gratings our system enables fully tailored holographic settings such as fringe density and shear, thus allowing flexible hologram demodulation, while maintaining simplicity and robustness. The performance is tested on a 3D-printed (using two-photon polymerization) brain phantom and fixed HeLa cells supplemented with cholesterol and oleic acids. Reconstruction analysis using the brain phantom indicates that the Hilbert method provides comparable results and resolution to the Fourier transform method in a significantly expanded measurement throughput. Our CP-HHT approach demonstrates the unique (not possible by fluorescence) high-throughput (especially when compared to cryogenic electron microscopy) capability to differentiate between cholesterol esters vs. triacylglycerol (TAG) rich lipid droplets (LDs), thus has potential for label-free biological research at sub-cellular level. The quantitative analysis of LDs’ refractive index emphasizes the method’s sensitivity in distinguishing between LDs with different neutral lipid content, offering new insights into LD heterogeneity, thus reinforcing the versatility and applicability of our CP-HHT system in broader bioimaging applications.

## Introduction

Quantitative phase microscopy (QPM) has emerged as a widely used tool for label-free imaging of cells and tissues. It leverages the inherent contrast produced as light navigates through transparent or semi-transparent samples, which is modulated by their intrinsic refractive index (RI) distribution and thickness. This principle has been pivotal in revealing invaluable insights across diverse fields, including biomedical sciences, flow cytometry^1^, neurosciences^2^, and cellular pathophysiology^3^. While conventional digital holographic microscopy systems, primarily based on the Mach-Zehnder interferometer setup^4^, have been at the forefront of QPM, they come with inherent challenges such as susceptibility to environmental perturbations and a need for high spatio-temporal coherence of illumination. The advent of common-path interferometric (CP-QPM) configurations has addressed some of these issues, offering resilience against environmental disturbances and greater flexibility in terms of light source coherence. Shearing systems offer the convenience of portability and ease of use; however, they only measure the derivative of the phase distribution in the shear direction, necessitating two orthogonal measurements to completely recover the object’s phase, typically through often numerical errors prone integration^5,6^. However, each CP-QPM variant, be it shearing^5^ or total shear system^7,8^, presents its own set of trade-offs. Consequently, shearing systems are best suited to objects with slowly varying phases due to their limited spatial resolution. Total shear systems, in contrast, tend to be cumbersome, require an extra 4f system, and depend on specialized, custom-made components such as specifically designed pinholes^9^.

To surmount these challenges, cutting-edge advancements like spatially multiplexed interferometric microscopes, which utilize diffraction gratings, have been introduced^7,10,11^. This evokes particular interest in holotomography (HT), where we have recently proposed a compact, Ronchi diffraction grating-based total-shear (coherence gating without the additional 4f system) CP-QPM system^12^. Notably, we also showed through simulations that diminishing the temporal coherence has a negligible impact on the 3D refractive index reconstruction quality^13^. However, Ronchi gratings have limitations, as they produce multiple diffraction orders with strong 0^th^ order, which adds both strong background and, in the case of coherent illumination, producing spurious spectral components. Additionally, the image processing for phase demodulation is traditionally limited to the Fourier transform method, which can be inefficient in terms of usage of the space-bandwidth product (SBP) of the optical system. Baek and colleagues^14^ proposed a sophisticated single-shot phase demodulation technique utilizing Kramers-Kronig relations (KK), which offers extended SBP in the frequency domain. This methodology facilitates the extraction of 2D phase information notwithstanding the complete overlap of the complex amplitude (coherent term) with its autocorrelation (incoherent) term within the frequency domain. However, using KK requires that the complex amplitude and its conjugate (both cross-correlation terms) are fully separated. This limitation can be overcome with the hologram demodulation using Hilbert transform, as we have shown in the previous works^15,16^. Consequently, in this contribution, we introduce a novel polarization gratings (PG) aided common-path Hilbert holotomography (CP-HHT) for high SBP tomographic imaging of refractive index distribution. This system employs a pair of PGs to allow for the adjustable fringe pattern composition and enables robust phase shifting through the use of a rotary linear polarizer.

Our novel system is experimentally tested with two kinds of samples. Firstly, we show results on the 3D printed brain phantom that was designed based on actual cuttlefish brain slices measurements. We believe that the proposed realistic phantom may be an interesting alternative to other 3D test objects for HT^17,18^. The brain phantom enabled verifying the great potential of Hilbert holotomography by emphasizing its capability to reconstruct fine object details. Furthermore, we propose application of Hilbert holotomography to an analysis of the 3D RI distribution of the lipid droplets (LDs) induced with cholesterol (CE) and oleic acids (OA) supplementation to cultured cells, showing, for the first time to the best of our knowledge, that RI tomography can successfully address aspect of LDs cellular heterogeneity in a label-free, noninvasive, high space-bandwidth fashion.

Understanding of LDs and their significant role in cellular processes has been enhanced by recent advancements in HT imaging of RI, through which profound insights have been provided and led to significant advancements across various biological studies^19^. The importance of 3D QPM for LDs analysis in live cells was highlighted, focusing on structural and biochemical parameters crucial for understanding metabolic diseases^20^. HT showed great usefulness for rapid lipid content quantification in microalgae, marking a step towards efficient biofuel production^21^. Hsieh et al. explored the regulation of LDs in response to fatty acid stimulation and starvation, providing metabolic disorder insights^22^. Integrating HT with fluorescence imaging for morphological and biophysical analysis of foam cells enabled assessing their therapeutic effects^23^, while the challenges of light scattering in HT were tackled by using non-toxic RI matching media, enhancing the fidelity of 3D RI reconstructions^24^. Low-coherence holotomography was used as a tool for long-term visualization and quantification of LDs in adipocytes^25^. A high-throughput comprehensive LDs visualization and characterization using a tomographic phase-contrast flow-cytometer was proposed by Pirone et al.^26^. Here we used the CP-HHT system for novel highly sensitive measurements of LDs RI distribution and differentiation of the cholesterol-rich (CE-enriched) vs triacylglycerol (TAG)-rich (OA-enriched) LDs, which showcases the type of LDs predominant in the analyzed sample. We used two sets of HeLa cells with LDs induced with cholesterol and oleic acids supplementation.

## Results

### Hilbert Holotomography

Hilbert spiral transform (HST), utilizing a background-free fringe pattern (hologram) and a fringe direction map, represents a significant advancement as a fully two-dimensional method over the one-dimensional KK technique. The implementation details of the HST phase retrieval algorithm are contained in Methods section. Here we focus on highlighting the merits of novel application of this phase retrieval approach in holotomography. With Fig. 1, we characterize the SBP conditions for phase demodulation using FT, HST and temporal phase shifting (TPS). Figure 1(a) shows the Fourier spectrum of the hologram with the high carrier frequency required for the FT method in the case of typical tomographic measurement. The pink circle denotes the filter size that is needed to exploit the entire NA of the optical system. Clearly, the variable oblique illumination, which is a vital element of HT, imposes a strong limitation on the SBP of the measurement (see the mutual displacement of the pink circles that corresponds to the illumination-related shift of the object terms in the hologram spectrum; here we consider the conical illumination scenario with fixed illumination tilt angle of 58° and variable azimuth). Thus, conventional single-shot phase imaging often resorts to using a smaller filter size for the FT phase retrieval, which numerically limits the spatial resolution. Another possible solution to bypass this problem is to increase magnification of the imaging system, which reduces the effective diameter (in a number of pixels) of the required filter and allows for increasing the fringe carrier due to enlarged Nyquist frequency. However, the increased magnification comes with the cost of reduced field of view (FoV). Thus, with both of these approaches, SBP of the measurement system is sacrificed. Our HST-based method, in contrast, is crafted to operate effectively even when the object’s frequency component (the first spectral order) cannot be completely separated within the Fourier space of the hologram and is overlapping with conjugate and autocorrelation terms (Fig. 1(b)). The proposed new CP-HHT approach therefore enables the efficient use of SBP of the measurement system by decreasing optical magnification, which enlarges FoV and thus increases the final result SBP. From comparison with Fig. 1(c) (hologram without the carrier frequency), it can be seen that HST provides comparable SBP conditions to TPS, and importantly, achieves it using a single frame per tomographic projection. Let us consider a practical example when the object under study has large SPB, i.e., it has large high spatial frequency content and at the same time is spatially extended (e.g., fragment of tissue or c. elegans bio-samples). Such an object is best imaged with microscope objective with high NA and relatively low transverse magnification (*M*) such as, e.g., water immersion objective with NA=1.2, 60˟. The described conditions result in challenging holograms with the spectral overlapping of object information, see Fig. 1 (d). In our example we assume the following realistic parameters of the HT system: camera pixel pitch *Δx* = 5 μm, light wavelength *λ* =0.5 μm, illumination incidence in air at 58⁰, an object immersed in an immersion oil with *n*_0_=1.515. It can be noticed that for all tomographic illumination angles the object term overlaps with the autocorrelation term and, for some illumination azimuths, also with the conjugated object term, prohibiting the usage of the FT method, while the application of HST phase retrieval is attainable.

**Figure 1.**
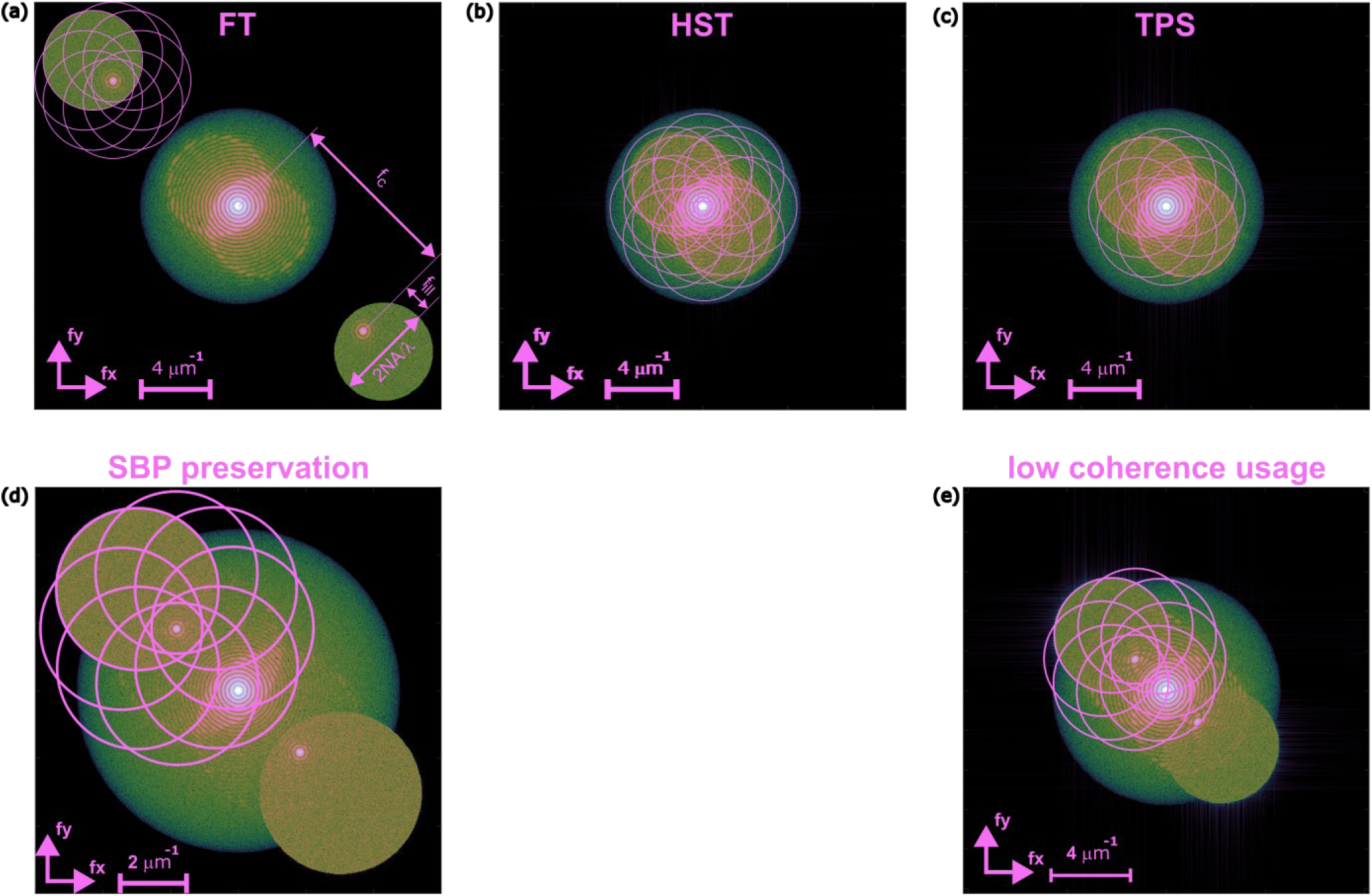
Comparison of the spatial frequency space conditions accessible for different methods of phase demodulation in holotomography: (a) FT, (b) HST, (c) TPS; symbols: *f*_c_ – fringe carrier frequency, *f*_ill_ = sinα_ill_/λ – illumination frequency; the graphs obtained for NA=1.45, λ=0.5 μm, illumination angle of 58° and conical illumination scenario*; two showcases of the HST relevance: (d) measurement SBP maximization, (e) application of the low coherence light sources*.

Besides the SPB motivation, we would also like to stress another important issue promoting the usage of HST phase retrieval in HT. By utilizing lowered coherence light source, we are able to reduce the coherent noise, which is fundamental for enhancing the final signal-to-noise ratio. It is particularly vital in phase demodulation due to the direct conversion of any intensity noise or spurious signals (e.g. parasitic interferences) into phase noise and, consecutively, refractive index reconstruction noise.

Notably, proposed CP-HHT system has the ability to demodulate the phase with decreased spatio-temporal coherence of the light source, as the interfering beams (+1 and –1 grating diffraction orders) travel exactly the same path, setting the optical path difference (OPD) to 0. Lowering the degree of coherence of light sources in interferometric systems has a major advantage of speckle noise and parasitic fringes reduction. However, with the reduced temporal coherence, the maximum OPD in the system has to be smaller than the coherence length *l_c_* of the source, which restricts the maximum possible carrier frequency of the interferometric image, potentially prohibiting the usage of the FT hologram reconstruction method.

To be more specific, let us assume a partially coherent light source with Gaussian shape of a spectrum, whose frequency bandwidth Δ*ν* is expressed with full width at half maximum. In this case the coherence length is given by:^13^

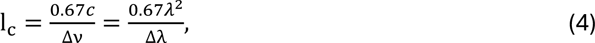

where *c* denotes the speed of light, *λ* is the light wavelength and *Δλ* expresses the wavelength spread of the source. For such a source, the maximum possible carrier fringe frequency that will not lead to the incoherence-induced drop of the fringe contrast is given by:

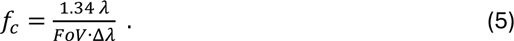

The problem of the numerical resolution limitation appears when the carrier frequency (Eq. (5)) is too small to ensure separation of the cross-correlationobjectterm and thus its full Fourier space filtering. For example, for the source with *λ*=0.5 μm, Δ*λ*=5 nm and *FoV*=100 μm, themaximumcarrierfrequency *f_c_*=1.34 μm^-1^. Fortheseconditionsand assuming the following typical tomographic parameters: NA = 1.45, *Δx* = 5 μm, illumination at 58⁰, *n*_0_=1.515, we obtain the hologram spectrum, see Fig. 1(e), with the evident spectral overlap that rules out the usage of the FT method. To achieve tomographic imaging in theseconditionsone may resort to the single frame HST phase retrieval method.

### Common-path Hilbert holotomography system

Figure 2 shows the experimental system used to obtain the hologram sequence and its further phase demodulation and 3D RI distribution reconstruction. It is conceptually driven by the open-top common-path grating based tomographic system^27^; however, we significantly advanced it by replacing the amplitude Ronchi grating with two PGs, which dramatically improves the performance and flexibility of the novel system. PGs are periodic structures that can interact differently with light depending on its polarization state.^28^ The grating lines are filled with liquid crystal molecules with changing orientation, so that they can manipulate light based on its polarization state. PGs are generating, for incident linearly polarized light, only two circularly polarized diffraction orders, +/-1^st^, with opposite handedness. On the other hand, when circularly polarized light is incident on the grating, the grating just deflects the light according to the deflection angle and rotated the handedness of the circular polarization (see Fig. 2c). This combination of two PGs and linear polarizer gives a lot of advantages over other well-established grating based common path systems: (i) there’s no 0^th^ order hence it will not affect the fringe pattern, (ii) the shear between both object beams can be easily adjusted with the grating-grating axial distance, (iii) both beams are traveling the same optical paths hence the system can be described as achromatic, (iv) fringe density can be tailored to the experiment conditions via the grating rotation (Fig. 2 c-d). This fourth property is the most interesting, as it allows for great system flexibility with very little operations – it can easily operate in the fully in-line configuration and perform the phase shifting with the rotation of the linear polarizer P (thank to the circular polarizations with opposite handedness), slightly off-axis and fully off-axis configuration. The on-axis and slightly off-axis configurations are the most advantageous in terms of the SBP of the system – we can utilize the full spatial frequency bandwidth and either focus on highest accuracy of the phase demodulation with TPS or marginally reduce the phase accuracy but increase the temporal resolution and use HST.

**Figure 2.**
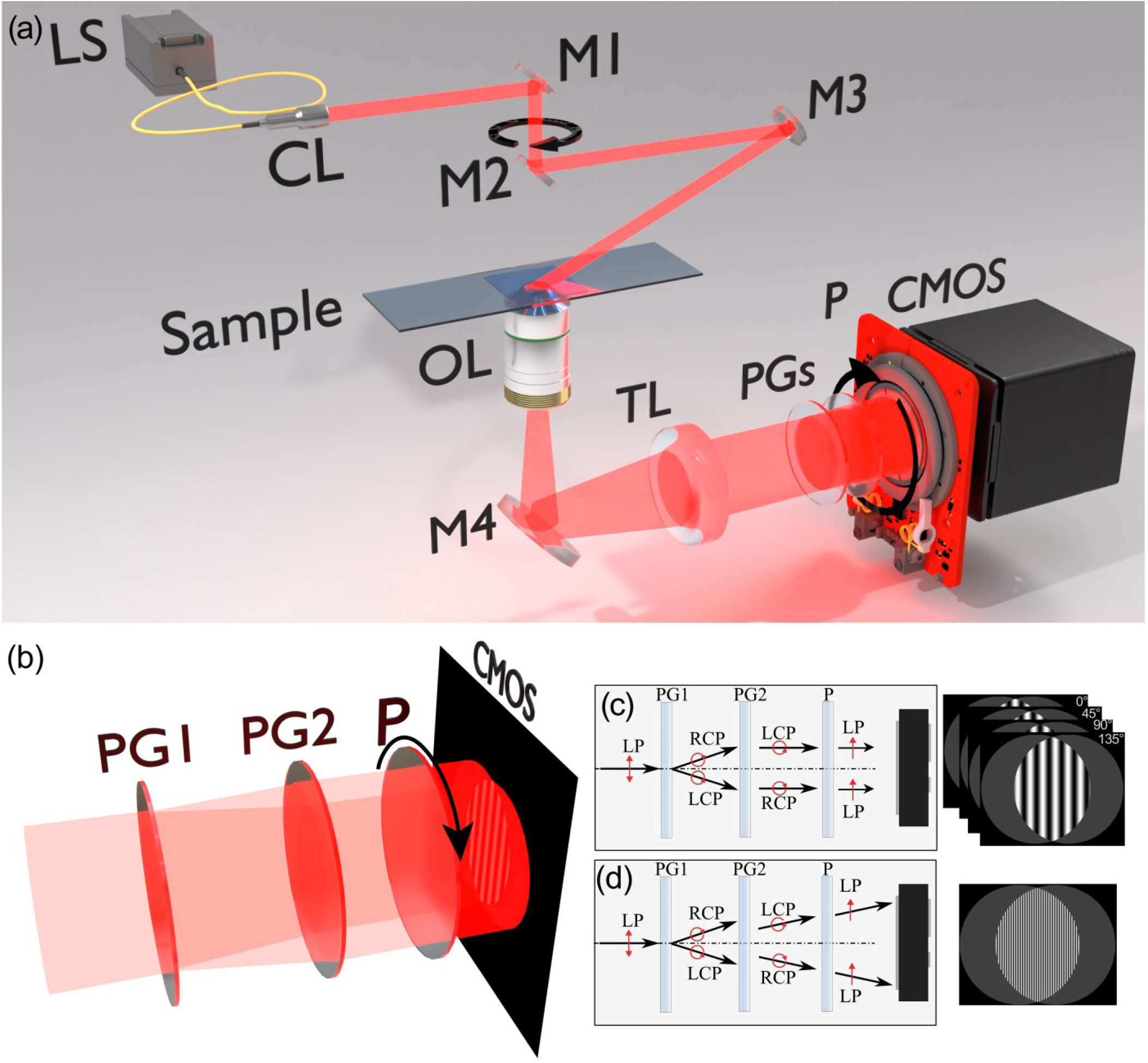
(a) 3D rendering of the proposed CP-HHT system: LS – light source, CL – collimating lens, M1-M4 – mirrors, OL – objective lens, TL – tube lens, PGs – polarization gratings, P – polarizer mounted on the rotation stage, CMOS – camera. (b) interference formation with PGs: (c) for the case of temporal phase shifting using the rotation of the polarizer, PG1 and PG2 lines are quasi-parallel to each other and images are acquired for 4 positions of the P: 0°, 45°, 90°, 135°; (d) for the case of single-shot HST with static P (0°) and slight inclination of the interfering diffraction orders, PG2 is slightly twisted with regard to PG1 to create the low carrier frequency.

Tomographic reconstruction was performed using direct inversion (DI) algorithm^29^ utilizing the first order Rytov approximation^30^. The theoretical foundations for DI were provided by Wolf within the generalized projection theorem^31^ that states that each 2D holographic projection of a 3D object carries information about a specific region, called Ewald cap, of the 3D object spectrum. DI is a straightforward implementation of this idea that applies direct interpolation of the object beam information in the spectral domain. We chose DI due to its popularity, open-access^32^ as well as robustness and fast computations. In our work, we apply the limited angle tomographic configuration that allows for convenient study of biological samples. This solution, however, suffers from the missing cone problem^33^ and thus decreased axial resolution of the tomographic imaging in comparison with the full angle tomography. Before tomographic reconstruction all holograms constituting the given tomographic data series were reconstructed (phase demodulated) with a hologram reconstruction method of choice. Then, the carrier frequency of the object beam was removed and the 2π phase discontinuities were eliminated with unwrapping algorithm^34^.

### Method validation with 3D printed brain phantom

We recorded a series of holograms for the set of various illumination angles. In our system we applied a conical illumination scenario where all illumination directions lie on a conical surface (Fig. 2a), the applied azimuthal step was 2°, which resulted in 180 projections in total. For TPS we recorded 4 phase shifted holograms for different orientations of the polarizer P (0°, 45°, 90° and 135°). These images were subsequently subjected to principal component analysis (PCA) based phase retrieval algorithm^35^ to reconstruct the complex optical field for each illumination direction. In the case of the HST method, only one fringe pattern image was captured for each illumination angle achieving quadruple acceleration of the recording process. Subsequently, these captured patterns underwent preprocessing, which involved denoising with the block-matching 3D filtering (BM3D) method ^36^ and background subtraction using the fringe pattern fast iterative filtering (fpFIF2) method ^37^. Following preprocessing, the complex optical fields were retrieved using HST.

As a validation object we used the custom made brain tissue phantom, an object that is a brain model of an adult female cuttlefish^38^. The phantom was fabricated using two-photon polymerization. Details of the phantom preparation are described in the Methods section. Figure 3 (a-c) shows the slice of the reconstructed RI distributions of the brain phantom using three hologram reconstruction methods and various fringe densities – (a) TPS for almost in-line configuration, (b) the HST method for low spatial carrier and (c) the FT method for same spatial carrier as in the HST method. Additionally, Videos S1-S6 show the corresponding 3D RI reconstructions using flythrough the stacks and maximum intensity projection for 3D visualization. We can clearly see from Fig. 3 that SBP for the HST-dedicated hologram (Fig. 3(b)) is significantly larger than for the FT-dedicated hologram (Fig. 3(c)), as the cross-correlation terms are overlapping one another, and FT method is significantly filtering the higher spatial frequencies for proper phase demodulation. Notably, HST method is also capable of achieving higher SBP compared to the KK phase demodulation method as the autocorrelation terms do not need to be separated from each other. The highest possible SBP is obtained through the TPS phase demodulation method, as the configuration can be quasi-inline, however, it requires 4 holograms. This method also provides the highest reconstruction quality; however, as can be noticed from comparison of Fig. 3(a), Fig. 3(b) and Fig. 3(c), the HST method provided comparable quality results, while enabling single-shot hologram registration (compared to the 4 holograms for TPS). This can be seen in the cross-sections of the brain features which are correctly reconstructed in both HST and TPS, while using FT method, due to decreased SBP, blurs higher spatial frequencies of the sample. Due to the inherent features of the HST method all important phase information can be transferred numerically after lowering the carrier frequency to fit the system’s SBP. HST method, in the case of physically too low SBP of the system for the FT method, still ensures efficient numerical transferring of all phase details in a single shot with the quality that is comparable to multi-frame phase shifting – please compare Figs. 3(a) and 3(b).

**Figure 3.**
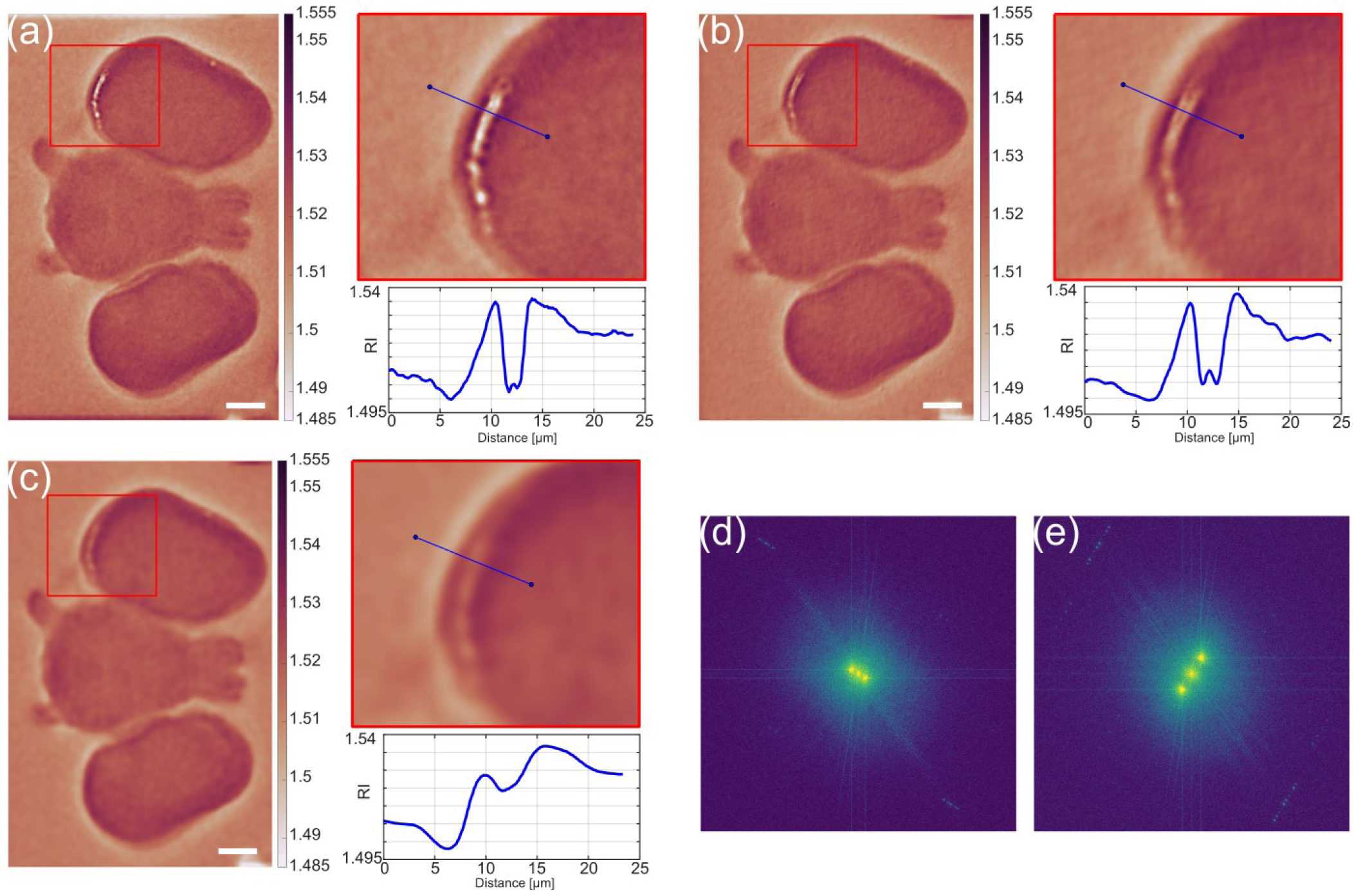
Comparison of a slice of the tomographic reconstruction using different hologram demodulation methods. (a-c) Tomographic reconstructions for (a) TPS, (b) HT and (c) FT methods; red rectangle shows the zoomed part of the brain and the blue line shows the cross-section position, graph shows the plot of the cross-section through the brain detail. (d-e) Fourier spectrum of a single hologram with fringe density for (d) TPS and (e) HST and FT. The scalebar is 10 µm.

### Quantitative analysis of the LDs

The goal is to validate if HT allows differentiating between cholesteryl ester-rich and triacylglycerol-rich LDs in a non-invasive fluorescence-free quantitative 3D high-throughput manner. LDs are crucial organelles for neutral lipid storage, with energy-rich triglycerides and sterol esters serving as lipid reserves essential for cellular balance ^39,40^. Keeping lipid levels balanced is critical for cell health. LDs, as key cellular organelles, manage lipid flows within cells. Altered LDs lipid composition or lipid composition in general are linked to pathologies such as atherosclerotic lesions^41^. Hence their precise analysis in terms of the lipid composition is of crucial importance and HT is a tool that can do it in a label-free quantitative way. We used HeLa cells with the increased either CE or TAG content (through cell supplementation with CE or OA, respectively).

We used the 650 nm wavelength to illuminate the sample and the illumination angle in air was 58°. Similarly, as with the imaging of brain phantom, we used 1.45 NA 100x objective lens and acquired 180 projections each with 4 phase shifted holograms. The holograms were reconstructed with TPS and submitted to tomographic reconstruction. In Fig. 4 we show the results of the imaging of LDs using proposed CP-HHT. Individual LDs are clearly visible, as their RI is significantly higher than the cytosol and other cellular organelles^19–21,25^. Following the measurements we carried out the segmentation for each of the LDs tomographic reconstruction. For quantitative LDs analysis, we used the following workflow: (1) segmentation of HeLa cells to localize LDs in the region of interest; (2) division of LDs into sub-volumes; (3) calculations of the mean RI value within the sub-volumes; (4) RI distribution analysis. Figure 4 shows the result of the segmentation of the analyzed LDs, (3D visualization of the tomographic reconstructions can be seen in Videos S7-S10).

**Figure 4.**
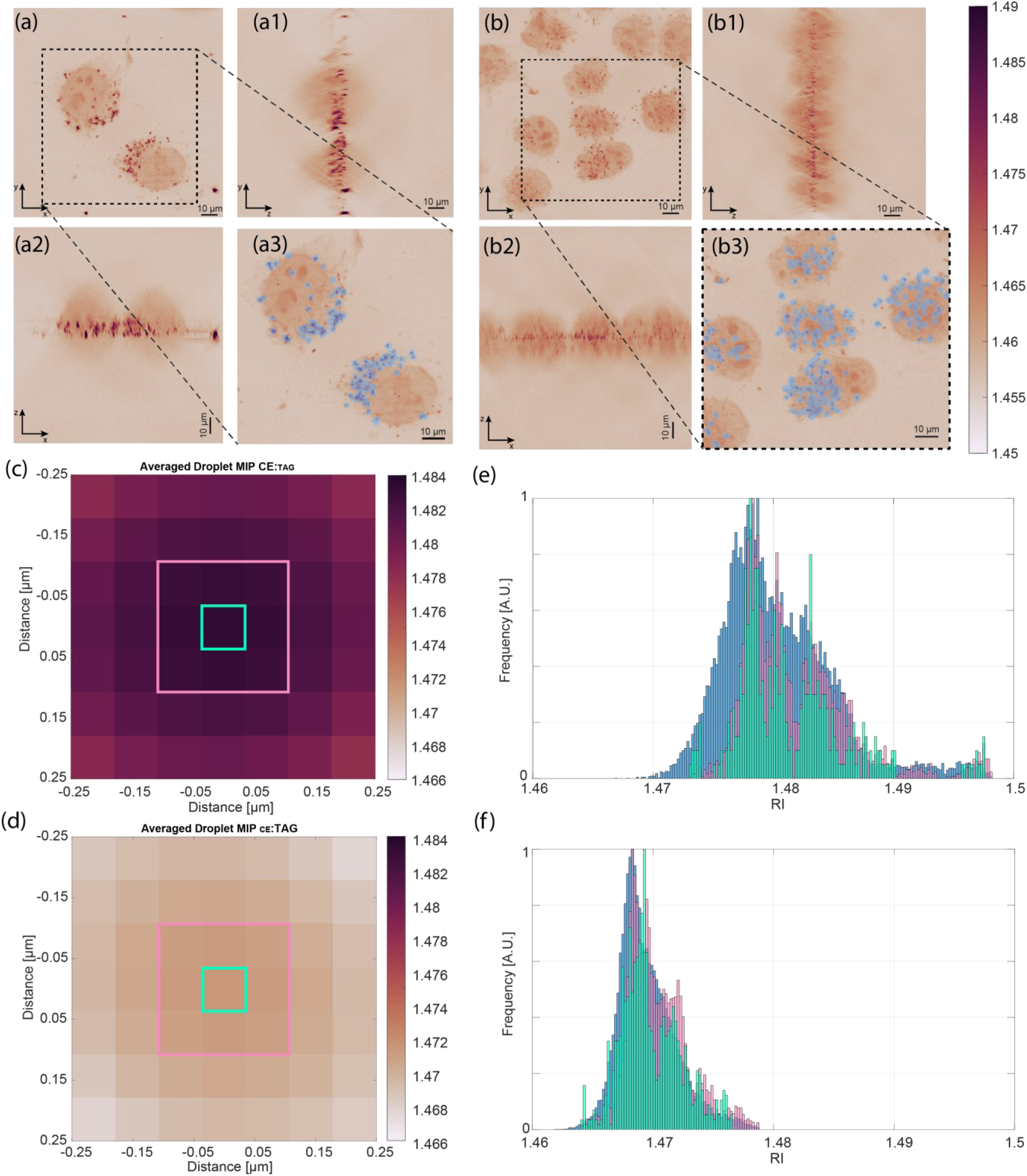
Holotomographic imaging and quantitative analysis of lipid droplets in HeLa cells. (a) Maximum intensity projection of the HeLa cell after **ON** treatment with **CE** with strongly visible LDs and its orthogonal views: (a1) YZ, (a2) XZ, and **closeup** (a3); (b) Maximum intensity projection of the HeLa cell after ON treatment with OA with strongly visible LDs and its orthogonal views: (b1) YZ, (b2) XZ, and closeup (b3); Averaged RI distribution of the LD for (c) CE and (d) OA induced LDs; (e-f) Histogram of the RI distribution for CE induced LDs (e) and OA induced LDs (f).

The distribution of the retrieved RI values for all analyzed sub-volumes is presented in Fig. 4 (c-f). Obtained histograms were calculated for three different subsets determined by the distance from the LD center (shown with colored rectangles in Fig. 4 c-d) to highlight the RI variations within LDs. Differences in the RI values between both types of LDs can be clearly observed. The OA-induced LDs exhibit a mean RI of 1.468, with a notably low standard deviation of 2.28 × 10^−3^, suggesting a more homogenous RI distribution, compared to the CE-induced LDs, which display a mean RI of 1.479 with a higher standard deviation of 4.91×10^−3^, which is an indicative of a more dispersed RI distribution. We believe that our analysis derives substantial support from the nuanced examination of RI distributions observed in HeLa cells samples. The RI distribution pattern is distinctively different for CE-induced LDs and its tomographic reconstruction shows that there are more RI values with significantly higher RI, meaning that the CE has strongly impacted the LDs.

These observations are quantitatively underpinned by the statistical analysis, which reveals significant deviations from normal distributions for both OA- and CE-induced cells (p-value < 0.001 for both groups and for both D’Agostino ^42^ and Anscombe-Glynn ^43^ tests), reinforcing the non-uniform nature of these distributions. Furthermore, Bartlett’s test indicated non-equal variances for both groups (p-value < 0.001), and Wilcoxon test indicated that the medians for both groups are not equal, providing additional certainty to the distinct characteristics of LDs induced by the two agents. The range of RI values also supports the notion of variability among CE-induced LDs (0.036) compared to OA-induced LDs (0.022), further underscoring the differences in distribution width and uniformity between the two groups. This analysis indicates differences in the lipid composition between the CE and OA induced LDs, which can be measured using our proposed high-SBP, robust and sensitive CP-HHT method.

## Discussion

In our study, we have developed and implemented a novel polarization grating-aided CP-HHT leveraging the extended SBP. The system stands out through its simplicity and robustness as it works in the common-path configuration. Thanks to the PGs, it can be easily adopted towards using different methods of hologram demodulation, including a straightforward implementation of TPS via rotation of the polarizer, without the need for implementation of piezoelectric stages. We showed that we can increase the SBP with single-shot phase demodulation using HST, offering results comparable in terms of quality to the ones obtained with TPS. This was verified with the 3D printed brain phantom samples. We applied the developed CP-HHT system for the RI analysis of two sets of lipid droplets in HeLa cells: induced with CE and OA. The proposed analysis indicates differences in the lipid composition between the CE and OA induced LDs, which can be measured using the proposed high-SBP, robust and sensitive CP-HHT method. Taken together, these measurements and statistical analyses showed distinctive differences in optical properties of LDs induced by OA and CE, with the latter showing an increased value of RI by approx. 0.01, suggesting an alteration in the lipid composition. This insight into the impact of supplementation substances on LD composition might help understanding their biophysical properties and functional roles within cellular contexts. Our study showed that holotomography can serve as a very powerful tool for label-free analysis of neutral lipid composition which could help uncover the role and occurrences of phase transitions of the lipids within LDs^39,41,44–47^. HT can complement data obtained by cryo electron microscopy (cryoEM) and other methods, especially since it is non-invasive, does not require meticulous sample preparation, is in orders of magnitude more cost-efficient, and most importantly, offers the possibility to image and measure living samples, which is a further step in understanding the influence and significance of cholesterol in LDs. While cryoEM provides high-resolution structural information and is invaluable for visualizing cellular components at the molecular level, it has several limitations when it comes to live cell applications. CryoEM requires complex and meticulous sample preparation, including rapid freezing and handling under cryogenic conditions to preserve the native state of the samples. This process not only adds to the cost and time but also restricts its application to fixed or frozen samples, making it impossible to observe dynamic biological processes in living cells. In contrast, HT enables the study of living cells without the need for any special sample preparation or staining, maintaining the natural physiological conditions. HT has the ability to perform live imaging (as the only limit in our case is the scanning speed, which can be significantly increased with the use of galvanometric scanners or DMD modulators) allowing researchers to monitor cellular processes in real-time, capturing dynamic events and interactions that would be missed in the static snapshots provided by cryoEM. Furthermore, HT’s high throughput capability makes it suitable for large-scale studies. Our proposed system has a potential to address question of LDs heterogeneity within the same cell culture, and with proper standardization might become a tool that will allow to estimate CE to TAG ratio based on RI of LDs enabling label-free high-throughput lipidomics with single LD precision.

## Methods

### Experimental System

Light source, LS, is a supercontinuum white light laser (NKT Photonics SuperK EVO), which is coupled into 100 µm core diameter multi-mode fiber. Then, at the output, the beam is collimated with the collimating lens CL and directed with mirrors M onto the sample. M2 and M3 are placed on the rotating arm generating oblique illumination incident on the sample. The angle of the illumination can be easily adjusted with the tilt of the mirror M3 and the height between the sample plane and the rotating arm. The sample is imaged on the CMOS sensor with objective lens OL (1.45NA 100x Nikon PlanApo) and tube lens TL. Between CMOS and TL we placed two PGs and a linear polarizer right before the camera (Teledyne Photometrics Prime BSI Express sCMOS camera), which is recording the resulting fringe pattern.

### Hilbert spiral transform phase demodulation

HST method applies a spiral phase function in Fourier space to produce a quadrature component relative to the fringe pattern under investigation generating complex analytic fringe pattern with easy access to the phase map. It is crucial to eliminate the noise and remove the background (incoherent) term while maintaining the fringe pattern’s cosine profile. Experimental (noise) and numerical (background) methods have been shown to yield promising results in this regard^15,37^.

For the sake of simplicity, if we disregard the noise, the input signal for HST, characterized as a background-free fringe pattern, can be described as follows:

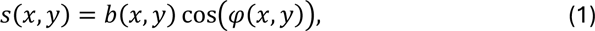

where *b*(*x*,*y*) and *φ*(*x*,*y*) represent amplitude and phase distribution, respectively. The output signal after HST is described as:

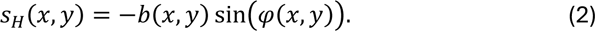

With the HST result, the phase distribution is a straightforward calculation:

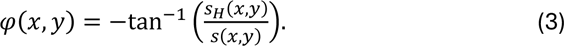

### Sample preparation

#### Brain phantom

We used the fabricated using two-photon polymerization system (Photonic Professional GT2, Nanoscribe GMBH). The original dataset comprises cross-sectional slices (100 µm thick) of the cuttlefish brain, which were stained using Phalloidin dye and imaged with Nikon AZ100 Multizoom Slide Scanner ^38,48–50^. We took the stack of 44 brain images and converted their grayscale intensities into laser power density values to achieve the desired RI distribution. This stack of images served as the design for fabricating a three-dimensional (3D) brain model using a two-photon polymerization setup. The technique enables to fabricate purely phase structures with the precise control over RI value, primarily determined by the material properties (in this case we used a polymer IP-Dip2 with a refractive index range of 1.530-1.547) and fabrication parameters (by adjusting the laser power density). Consequently, our approach facilitated the fabrication of a 3D brain model with phase-encoded histological features.

#### HeLa cells

LDs in control HeLa cells were cultured in DMEM GlutaMAX media (4.5g/L D-Glucose, Penicillin, Streptomycin and 10% FBS) consisting a mixture of cholesterol esters (CE) and triacylglycerol (TAG). HeLa cells were seeded on 3 cm, glass bottom cell culture dish in Dulbecco’s Modified Eagle Medium (DMEM). To induce incorporation of CE into LDs, one set of samples was over night (ON) supplemented with 25 µM cholesterol (CE, RI=1.47±0.06^51^), while to trigger the incorporation of TAG, supplemented ON with 200 µM oleic acids (OA, RI=1.459^52^). Both types of HeLa cells were fixed with 4% paraformaldehyde solution for 30 min on the imaging grade glass bottom Petri dishes (Greiner, cat nr 627860) and submerged in phosphate-buffered saline (PBS) for imaging.

## Supplementary material

See Visualizations 1-8 for supporting content.

## Funding

National Science Center, Poland (2020/37/B/ST7/03629). Warsaw University of Technology within the Excellence Initiative: Research University (IDUB) programme. WD was supported by the EMBL

Interdisciplinary Postdoctoral Program (EIPOD4) under the Marie Skłodowska-Curie Actions COFUND. The research was carried out on devices cofunded by the Warsaw University of Technology within the Excellence Initiative: Research University (IDUB) programme.

## Supporting information

Visualisation 1

Visualisation 2

Visualisation 3

Visualisation 4

Visualisation 5

Visualisation 6

Visualisation 7

Visualisation 8

Visualisation 9

Visualisation 10

## Acknowledgments

We would like to thank Julia Mahamid for insightful comments.

## Data availability

Data underlying the results presented in this paper are not publicly available at this time but may be obtained from the authors upon reasonable request.

## Author information

P.Z conceived the idea of using polarization gratings and Hilbert transform in common path holotomography system, designed and built the optical system, performed experiments, developed the acquisition pipelines, and analyzed the data, J.W. performed theoretical analysis and simulations, provided the algorithms and carried out the tomographic reconstructions, M.R. provided image processing tools, M.J.M. performed the data acquisition and tomographic reconstructions of the LD samples, and analyzed the LDs data, E.W. fabricated the brain phantom sample, W.D. prepared the LD samples and provided the idea of their measurements, M.J. helped with the optical system design and development, M.T. supervised and acquired funding for the project. P.Z. wrote the manuscript with contributions from all authors.

## Notes

### Competing Interest Statement

The authors have declared no competing interest.

### Summary of Updates

Figure 1, Figure 3 and their respective descriptions

